# Distinct yet neighboring neural populations encode past, future, and surrounding speech context in the human temporal lobe

**DOI:** 10.64898/2026.05.13.724774

**Authors:** Marianne de Heer Kloots, Atlas Kazemian, William Turner, Josef Parvizi, Laura Gwilliams

## Abstract

Context is critical for both human and artificial speech comprehension systems. While the role of preceding context in speech processing has been well documented, the neural mechanisms supporting the integration of subsequent input — phonemes and words that occur in the future — remain poorly understood. Here, we leverage advances in artificial speech systems to model the contribution of different sources of context on the neural encoding of speech in the human brain. For neural encoding, context-informed but not context-uninformed speech model embeddings explain unique variance in human neural activity beyond acoustics, including in early speech processing regions. In particular, model embeddings informed by past, future, and surrounding context explain activity in distinct intracranial electrodes. These electrodes are left-lateralised, and spatially intermixed in the temporal lobe. We find that beyond-word context is crucial for the representational quality of speech model embeddings, and in particular for the encoding of abstract linguistic information. Our finding that spatially neighboring yet distinct neural populations in the temporal lobe encode representations shaped by different contextual sources (past, future, and surrounding input) provides key insight into the neural circuitry that integrates multiple forms of contextual information. Furthermore, our results may inform the downstream use of self-supervised speech representations in language technology tasks, and in models of speech comprehension in the human brain.

## 1 Introduction

Language understanding involves continuously integration of incoming input with available contextual information. Better modelling of context has been one of the major drivers behind recent progress in language technology, allowing both text- and speech-based systems to better capture input data structures and flexibly adapt to task demands (Brown et al., 2020; Omnilingual ASR team et al., 2025; Roll et al., 2025; Lampinen et al., 2025). At the same time, the use of contextualized predictions has been proposed as a key factor of alignment between human and artificial language processing, leading to the success of artificial language systems in predicting human brain activity (Jain and Huth, 2018; Schrimpf et al., 2021). In line with this, model-brain alignment tends to improve with more prior context provided to the model (Abnar et al., 2019; Anderson et al., 2024).

The human brain indeed uses contextualized predictions in processing speech. Information across timescales is integrated to form multi-level predictions of the future (Heilbron et al., 2022; Gwilliams et al., 2025b), and contextual cues (e.g. word, sentence, and speaker information) can affect the perception and recognition of linguistic units such as phonemes or words (Marslen-Wilson, 1984).

Importantly, context effects in human speech perception are not only characterized by the length of available context, but also by its *type* (future or past). Both neural and behavioural evidence indicates that the human brain informs its representations of spoken language input through a combination of *prediction* (using preceding context to anticipate upcoming input; (Kuperberg and Jaeger, 2016; Federmeier, 2007)) and *postdiction* (using subsequent context to shape representations of prior input; Ganong, 1980; Connine et al., 1991; Gwilliams et al., 2018). Hence, human contextualized language processing seems driven by flexible mechanisms, which extract information from different sources of context, rather than by solely maximizing the amount of integrated prior content.

How do human and artificial neural networks integrate contextual cues during speech processing? Here we take first steps in answering this question, by studying contextualized word representations extracted from a self-supervised speech model (Wav2Vec2; Baevski et al., 2020). Specifically, we explore how the model’s representations of words are affected when manipulating *what context is available* beyond the target word (see Figure 1); both in terms of **context size** (number of context words) and in terms of **context type** (past, future, or surrounding context). We first analyse how these context manipulations affect the model’s representation of speech content from a spoken narrative, and find that beyond-word context is crucial for the model’s extraction of abstract (linguistic) word-level features from its acoustic input. We then use this model and its context-manipulated representations as a tool to understand the encoding of past, future, and surrounding context in human cortical activity (Figure 1). All three context types substantially improve the prediction of neural representations beyond acoustic features and isolated word embeddings. Moreover, this improvement manifests within a short time window after target word onset (i.e. during the brain’s encoding of the *target word*, rather than its encoding of the context itself), and we can identify separable neural populations which are better modelled with either past, future, or surrounding context. These findings highlight the exciting potential of using self-supervised speech model representations to improve our understanding of speech contextualization in the brain.

**Figure 1:**
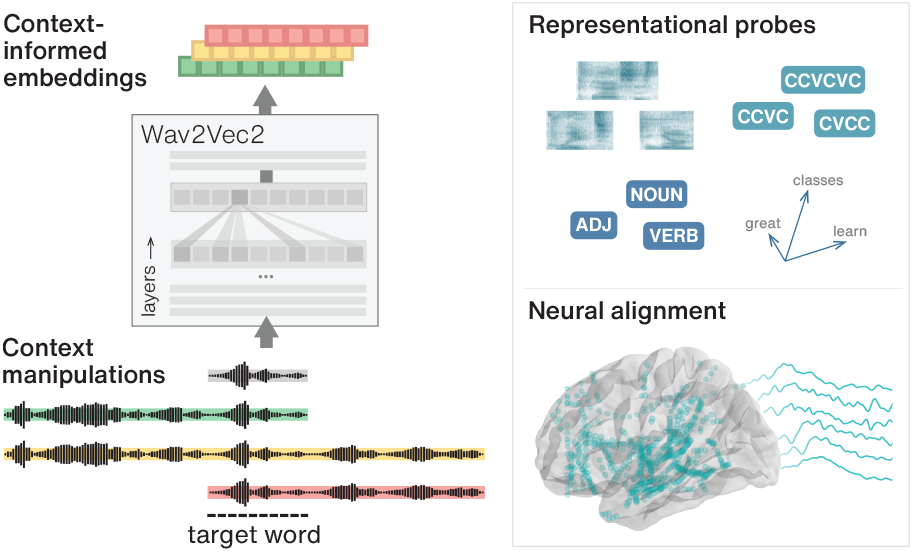
We analyze effects on Wav2Vec2’s encoding of word-level information as well as its alignment to human neural activity recorded with stereo-electroencephalography, when manipulating both the *size* and *type* (preceding, surrounding, future) of beyond-word context available at model input.

## 2 Related work

### 2.1 Neural representational alignment in language processing

Representations from speech- and text-based language processing systems show substantial alignment to human neural activity. The reason for this alignment, however, remains debated. While some take the superior neural predictivity of autoregressive language models as evidence that models and brains employ *shared objectives* for next-token prediction (Schrimpf et al., 2021; Goldstein et al., 2022; Caucheteux et al., 2023), it’s possible that the observed results are instead driven by inherent co-occurrence statistics in natural language data (Schönmann et al., 2025), or by the general richness of model-internal representations generalizing to many tasks beyond next-token prediction (Antonello and Huth, 2024). Another view points to geometric signatures of brain-aligned model representations, which follow the same layerwise pattern as the decodability of more abstract (but not surface-level) linguistic information — hence, model-brain alignment may be driven by *shared abstractions* from stimulus input (Cheng et al., 2026), rather than shared objectives or mechanisms.

Effects of context on representational alignment have so far mostly been studied in *text-based* language models. In text-based language models, the integration of prior context shapes both the representational geometry of model-internal activations (Hosseini and Fedorenko, 2023) and their neural alignment (Jain and Huth, 2018; Toneva and Wehbe, 2019), including in early auditory regions (Mischler et al., 2024). Interestingly, alignment to human language processing measurements is not necessarily optimized with maximal context: prediction of reading times as well as fMRI signals is in fact optimized by *restricting* access to limited amounts of prior context (Kuribayashi et al., 2022; Tikochinski et al., 2025).

Representations extracted from textual inputs nevertheless remain fundamentally limited for modelling neural activity involved in speech comprehension. *Speech-based* models operating on audio inputs offer an interesting alternative; specifically, the layerwise neural predictivity of self-supervised speech model (S3M) representations follows the cortical hierarchy of speech processing (Vaidya et al., 2022; Millet et al., 2022; Li et al., 2023), and the areas and features driving S3M alignment to fMRI recordings have been shown to be distinct from those of text-based models (Oota et al., 2024). Moreover, speech-based models show mechanistic alignment to human neural activity in their encoding of linguistic information: the neural dynamics of word-form encoding in auditory cortex align with those in S3M representations (Zhang et al., 2026). Given these findings and the contextualized nature of speech model representations, it is likely that word-level information (potentially shaped by beyond-word context) could play a substantial role in S3M-brain alignment, though not yet systematically investigated in existing work.

### 2.2 Context and linguistic structure in S3M representations

Modern S3M architectures for speech representation learning are trained on speech recording segments of around 20 seconds in length (e.g. Baevski et al., 2020; Hsu et al., 2021; Chen et al., 2022). Raw waveforms are internally transformed by a convolutional feature encoder into sequences of frame representations with a receptive field of 25 ms. A Transformer-based contextualization module subsequently integrates information across frames using a bidirectional attention mechanism, such that output frame representations can be informed by any other frame in the input.

In practice, a variety of local and mixed attention patterns emerge across model-internal layers, some of which are aligned with linguistic units such as phones (Shams and Carson-Berndsen, 2024). Sensitivity to local phonological context has been observed in early hidden-layer representations (de Heer Kloots and Zuidema, 2024; Gessinger et al., 2026), while word-level information tends to peak in middle-to-late layers (Pasad et al., 2024; Sauter et al., 2026). Measurements of effective context use have shown that greater contextualization is correlated with improved word-level task performance (Meng et al., 2025), and hidden-layer representations are significantly affected by how much beyond-word context is provided at input (Choi et al., 2024). Furthermore, transcription-finetuned models have been demonstrated to use non-local lexical information in disambiguating homophone orthography (Mohebbi et al., 2023).

In sum, both human biological and artificial neural networks appear to make use of context-informed word representations in speech processing. However, to our knowledge, how variations in context size and type shape word-level information represented in speech models, and how such context variations affect model-to-brain alignment, has not been systematically investigated. These two questions form the focus of our current study.

## 3 Data

### 3.1 Speech recording

For all analyses in our study, we make use of a 15 minute spoken narrative recording for which neural data of 10 listening participants has been collected (see section 5.1). The recorded narrative is a public speech by a single male speaker covering lessons learned in his career and personal life. The narrative was automatically transcribed and force-aligned to text on the word level using Whisper-large-v3 (Radford et al., 2022), after which errors were manually corrected. The input for all our analyses is sampled from a total of 2001 word occurrences across the narrative, with durations ranging between 30 and 1000 ms (mean duration: 260 ms).

### 3.2 Transformer model embeddings

To obtain Wav2Vec2 embeddings for analysis, we feed the spoken narrative as input to the model. We extract CNN output, input embeddings and layerwise hidden state representations from the original Wav2Vec2-base model pre-trained on 960 hours of audiobook recordings (Baevski et al., 2020), as well as a randomly initialized untrained model with the same architecture, for comparison. To obtain word embeddings from each layer, we follow earlier work analyzing word-level information in S3M models (Pasad et al., 2024) and mean-pool across audio frames within each word. We use a sliding window across words to obtain embeddings for each target word, while varying the amount of preceding and future words included in model input to manipulate the length and type of context available to the model to inform its representation of the target word (see Figure 1). To ensure that we measure effects of included speech context and not model input duration, we pad all input audio segments to be 20 seconds in duration, by adding silence around the speech content (word+context) for each input given to the model. In total, we extract embeddings for isolated words as well as 20 context-informed conditions, varying context type (preceding, surrounding, and future) and length (between 1 and 20 words).

## 4 Context effects on model-internal representation structure

### 4.1 Layerwise scoring measures

To explore how the availability of different amounts and types of context affects the representation of words in Wav2Vec2, we use four diagnostic probes to analyse how different kinds of information are represented in the extracted embeddings, ranging from acoustic to higher-level linguistic features. Given the limited amount of narrative embeddings to train and evaluate probes on, we use a combination of two relatively light-weight analysis techniques to compare model embedding structure to interpretable feature spaces and categories: Canonical Correlation Analysis (CCA; Hardoon et al., 2004) and Linear Discriminant Analysis (LDA; Tharwat et al., 2017). Variants of CCA and LDA have both been used in interpretability analyses of word representations in speech models before (Pasad et al., 2024; de Heer Kloots et al., 2025, 2026). We use CCA to quantify alignment to continuous features (MFCC features for *acoustic alignment* and GloVe features for *semantic alignment*), and LDA to quantify clustering according to structural categories (word form clustering by syllable type structure, and syntactic clustering by part-of-speech category). *CCA scores* are the mean Pearson’s correlation between the CCA components projected from each model and target feature space; *LDA scores* are the mean silhouette scores measuring cluster separability in LDA-projected space; for both, more positive scores indicate better alignment and scores of 0 correspond to random structure. We evaluate the layerwise probes across 5 independent train/test splits sampled from subsets of data samples designed for each probe, with constraints on splits to avoid analysis confounds (further details are included in Appendix A).

### 4.2 Projection similarity comparisons

Both CCA and LDA scores are computed after linearly projecting from model-representation space to lower-dimensional subspaces, which are optimized for alignment with the target feature space (MFCC- and GloVe-CCA) or for separability of the target categories (SylType- and POS-LDA). This means that both CCA and LDA can additionally serve as supervised dimensionality reduction techniques, where the dimensionality-reduced projections preserve any information in model representations that helps alignment to target features. After computing our probing scores on the CCA- and LDA-projected subspaces, we use them to quantify how *similar* the feature-tuned subspaces are across all context conditions. We quantify similarity between projection spaces using Representational Similarity Analysis (Kriegeskorte et al., 2008), constructing cosine-based dissimilarity matrices within each projection space and visualizing Pearson’s correlation scores between them as a measure of similarity.

### 4.3 Results

We visualize the results of our representational analyses across context conditions in Figure 2. We observe that increasing context *size* generally improves Wav2Vec2’s encoding of all interpretable feature structures we probed. This highlights the importance of including beyond-word context for the representational quality of Wav2Vec2 embeddings, even for the encoding quality of relatively low-level acoustic structure. Interestingly, for any level of information beyond acoustics, isolated word embedding probes hardly reach any above-random clustering or alignment scores. In line with this, probing results for isolated word embeddings (i.e. the no context condition) show no difference between trained and untrained Wav2Vec2 models (Appendix A.5). This means that including beyond-word context is crucial for any kind of word-level linguistic structuring to get encoded in Wav2Vec2 representations of spoken narrative data.

**Figure 2:**
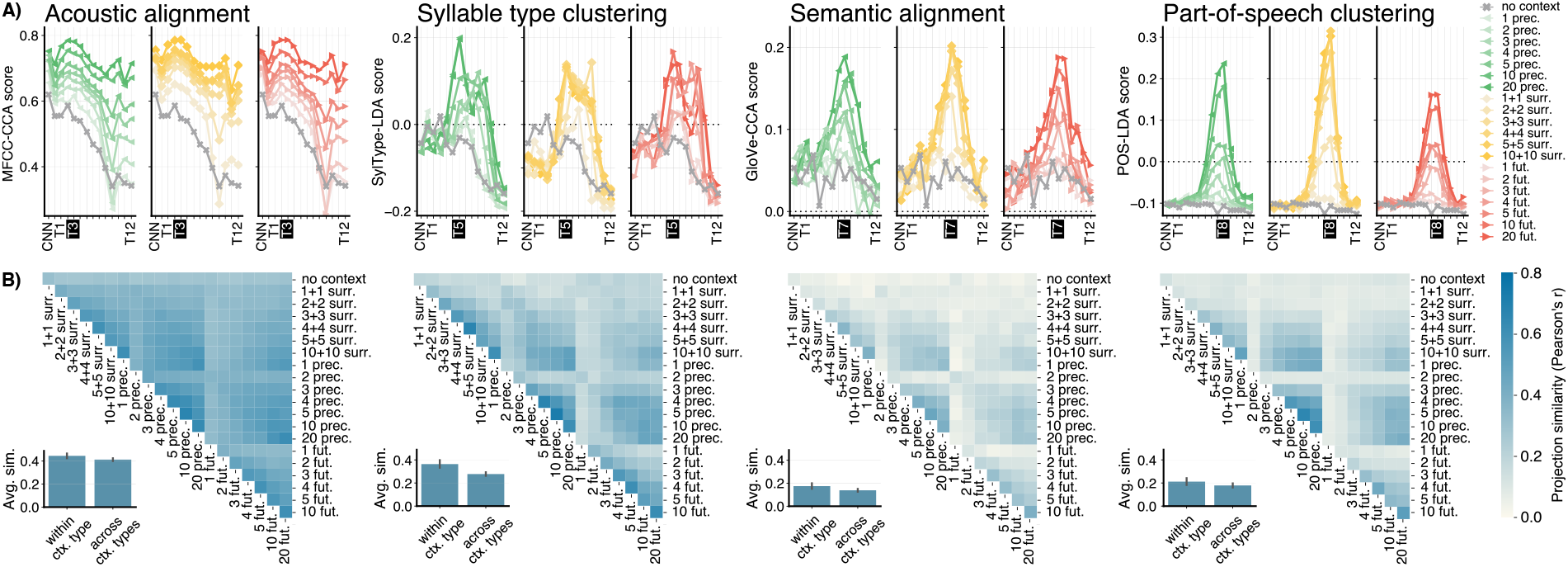
Context shapes the encoding of words in Wav2Vec2: at all levels of analysis, information about the target word is better represented when processed in context vs. in isolation. A) Layerwise results for each context condition across 4 representational probing tasks. Scores are on different scales across tasks and only displayed together to demonstrate the variety in relative layerwise peaks; black dotted lines indicate random structure. B) Projection similarities between context conditions for the top layer of each probe (highlighted in black).

Among context-informed embeddings, we observe that each context type does support the encoding of both acoustic and linguistic structure, with higher-order linguistic information (as measured by GloVe-alignment and POS-clustering) generally peaking in higher Transformer layers (7-8) than word form structure (layer 5) and acoustics (layer 1-3). Between the three context *types*, probing results mostly show little difference in the encoding of interpretable structures, except for part-of-speech clustering. This suggests that context *type* may primarily affect Wav2Vec2’s encoding of word properties for which neighboring word context could be crucial in achieving the correct interpretation (such as determining a word’s part-of-speech category).

Finally, we compare the similarity of extracted information between probes for all context conditions, as measured by the representational similarities between the probes’ fitted projection spaces. We observe that similarity is higher between embedding conditions with more shared context. Overall similarity for embeddings extracted with little-to-no context (i.e. one or two context words) is low. Across levels of information, we see that projection similarities are highest between the acoustic projections, and lower for probes decoding more abstract linguistic structuring. In general, projection similarity patterns follow those of our layer-wise scoring metrics: with improved alignment to interpretable feature structures, probe projections across different context conditions are more aligned with each other, as well as with the probed target feature spaces. However, despite seeing little effect of context type on layer-wise scoring metrics, we do find projection similarities are usually greater within than across context types, potentially indicating that while each context type enables similarly accurate linguistic structuring, there are qualitative differences in the information that different types provide to support such structure.

## 5 Context effects on neural alignment

### 5.1 Neural data

To compare model representations to human neural activity in speech comprehension, we make use of *intracranial stereo-EEG* (iEEG) data collected from patients receiving treatment from the Stanford Epilepsy Center, who volunteered to participate in data collection after having iEEG depth electrodes implanted as part of their clinical care. Participants listened to a series of audiobook snippets while neural activity was recorded. Participants reported no difficulties in language comprehension. We aggregated iEEG data recorded during narrative listening across a total of 1028 electrodes in 10 participants. Recorded electrode data was preprocessed using standard techniques for intracranial data analysis: the neural signal was aligned to the speech stimulus, and band-pass filtered to extract high-gamma activity (70-150 Hz). Word-locked epochs were then extracted between 500 ms before and 1500 after word onset, and epoch activity was down-sampled to 100 Hz.

For our electrode-level neural similarity analyses, we divided electrode signals into 20 time windows across the epoch (each spanning 100 ms and containing 10 samples of subsequent neural activity). These time-windowed neural activity vectors form the basis of the iEEG-based representation space for our neural similarity analyses.

### 5.2 Representational similarity analyses

We are interested in alignments between Wav2Vec2 and human neural representation spaces, but in particular in alignments driven by higher-level context-informed features, abstracting away from local acoustic properties in the speech signal.

Our approach to neural-representational modelling therefore follows earlier work in the visual domain, employing a version of Representational Similarity Analysis (RSA; Kriegeskorte et al., 2008) that uses cross-validated non-negative least squares (NNLS) to model the neural dissimilarity space as a weighted sum of feature dissimilarity spaces (Khaligh-Razavi and Kriegeskorte, 2014; Jozwik et al., 2016). To control for acoustic effects on similarity between Wav2Vec2 and iEEG representations, we include two acoustic feature spaces in our representational model, and analyze our effects of interest on the *unique variance* explained specifically by the Wav2Vec2 features, beyond acoustic correlates (see Figure 3).

**Figure 3:**
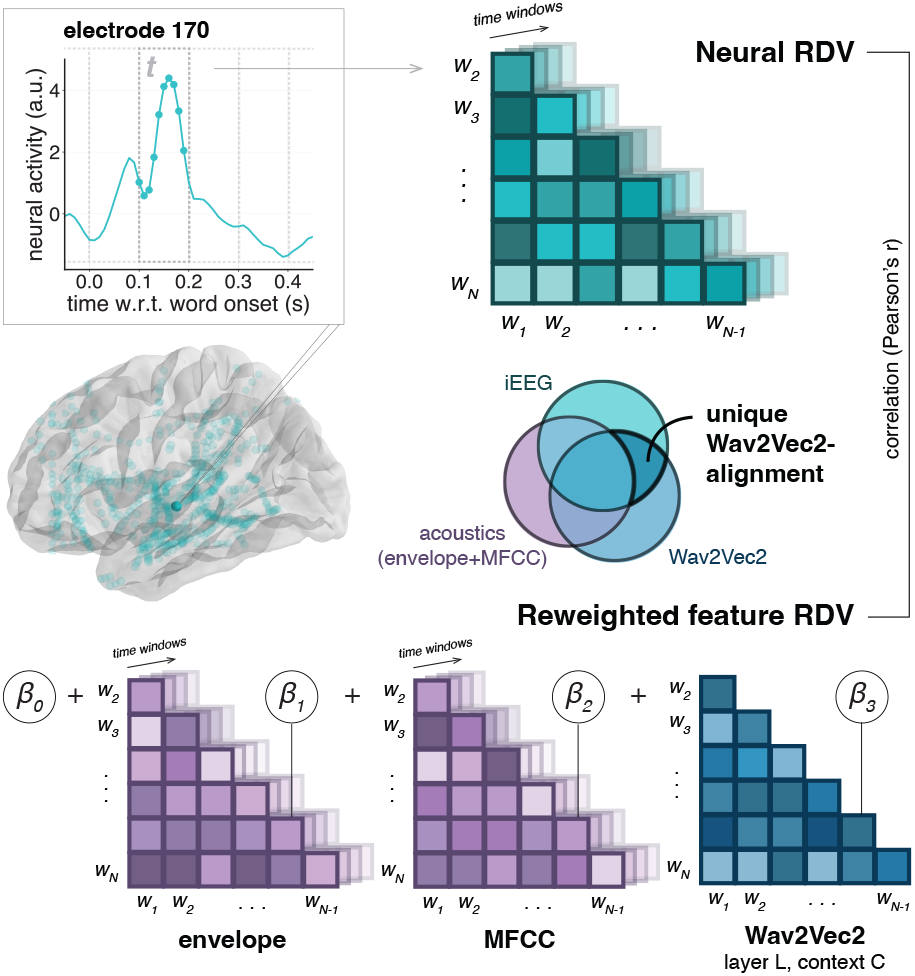
We compute Representational Dissimilarity Vectors (RDVs) for word-locked iEEG data, the speech envelope, and MFCC features (all at each epoch time window), and word embeddings extracted from Wav2Vec2 (at each layer and context condition). Our representational model is a weighted sum over feature RDVs; scalar weights *β* are optimized to predict the neural RDV. We measure unique Wav2Vec2-alignment as the relative increase in correlation between the feature-reweighted RDV and neural RDV, when including Wav2Vec2 vs. only acoustic features.

Across three folds, we fit a separate NNLS model over feature RDVs for each epoch time window, model layer, and context condition, optimizing weights on two-thirds of the narrative recording; we then compute the model’s *RSA score* as the Pearson’s correlation of the reweighted feature RDV and the neural RDV on the held out third. The unique variance explained by Wav2Vec2 is the difference in RSA score for an acoustics-only (envelope+MFCC) vs. full (envelope+MFCC+Wav2Vec2) feature model. We further compute a *context-advantage* score for each context-informed embedding condition, defined as benefit to unique Wav2Vec2-alignment compared to the context-uninformed condition, and a *context preference* score, quantifying the relative benefit of each context type. More details on the structure of our representational model, its evaluation and these scores are included in Appendix B.

### 5.3 Results

We computed acoustics-only (envelope+MFCC) as well as full model RSA scores for each epoch time window across all 1028 electrodes. Observing the general pattern of results, we see that both acoustics-only and full model scores are highest in auditory and early language processing regions, across both hemispheres (Figure 4, Appendix B). As we are primarily interested in analyzing variation in Wav2Vec2-alignment for electrodes with enough signal to be modelled well, we choose to further analyze a subset of electrodes based on a minimal threshold of full model RSA scores (0.0015) – this results in a selection of 355 electrodes where RSA scores exceed the threshold for at least half of the time windows in the epoch, across all three folds used for cross-validation. Additionally, we focus our analyses on scores observed for Wav2Vec2’s 7th Transformer layer, where relative Wav2Vec2-alignment compared to acoustics-only scores peaks (Appendix B).

**Figure 4:**
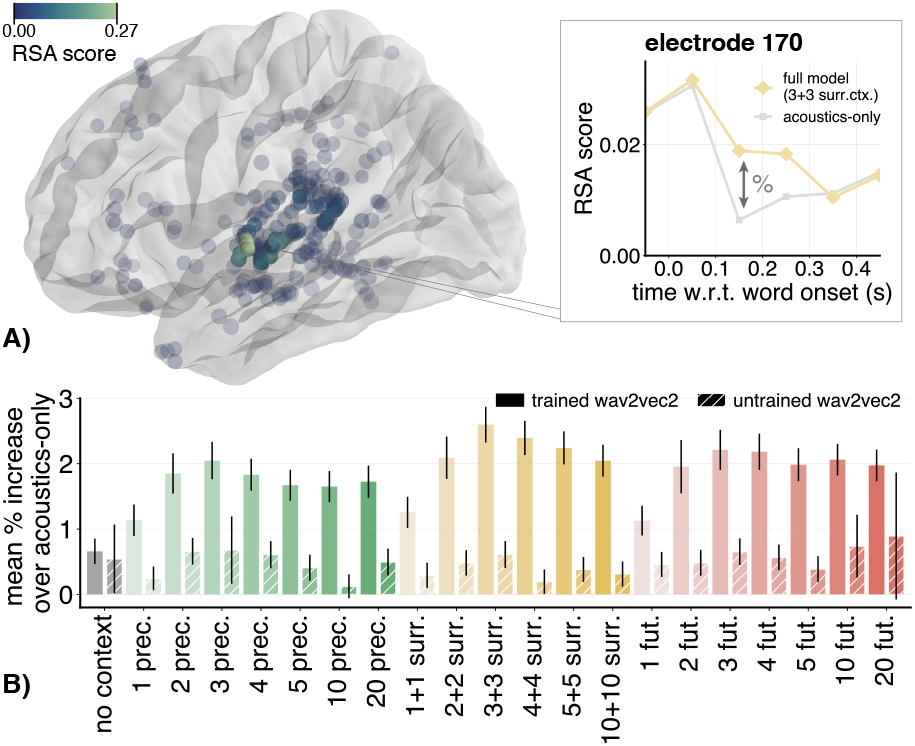
Context greatly improves the neural alignment of Wav2Vec2 embeddings to word-locked iEEG data. A) Full model RSA scores are highest around auditory processing regions, but electrodes do show substantial Wav2Vec2-alignment beyond low-level acoustics. B) Context benefits in Wav2Vec2-alignment do not arise in embeddings extracted from an untrained model, and show variation across both context sizes and types.

Comparing the acoustics-only and full model results (Appendix B), a large amount of variance in neural representation space for these electrodes can be attributed to low-level acoustic features. Nevertheless, Wav2Vec2 embeddings do substantially improve neural alignment beyond low-level acoustics, for many electrodes spread across various anatomical regions (Appendix B).

We next test whether context benefits the neural alignment of Wav2Vec2 embeddings. Averaging relative Wav2Vec2-alignment scores for each context condition across the epoch, we observe that including up to three words of preceding and/or future context substantially improves the alignment between word representations encoded by Wav2Vec2 and word-locked iEEG activity patterns (Figure 4). Furthermore, when comparing across context types, we see a small benefit for surrounding context embeddings as compared to embeddings with access only to preceding or future context. This could indicate that Wav2Vec2 effectively uses up to 3 bidirectionally surrounding words to inform the aspects of its word representations that increase alignment with human neural activity. Remarkably, Wav2Vec2-alignment for the no context condition is much worse than for the context-informed conditions, with similar scores obtained for the trained and untrained model embeddings – this mirrors our representational probing analyses. Finally, we explore how context-advantage patterns vary across the neural populations measured by each electrode. While we saw slight differences between context types when aggregating data across all electrodes and epoch time windows (Figure 4), it is not obvious whether this result arises because the degree of neural alignment follows a similar pattern across context conditions and/or time windows in all electrodes, or whether there are electrode-wise differences in which context condition best models their activity.

Figure 5 demonstrates that we observe the latter: some neural populations across the language network preferentially encode a single context type, while other electrodes encode multiple context types; either simultaneously (e.g., highlighted electrode c), or sequentially (highlighted electrode e).

**Figure 5:**
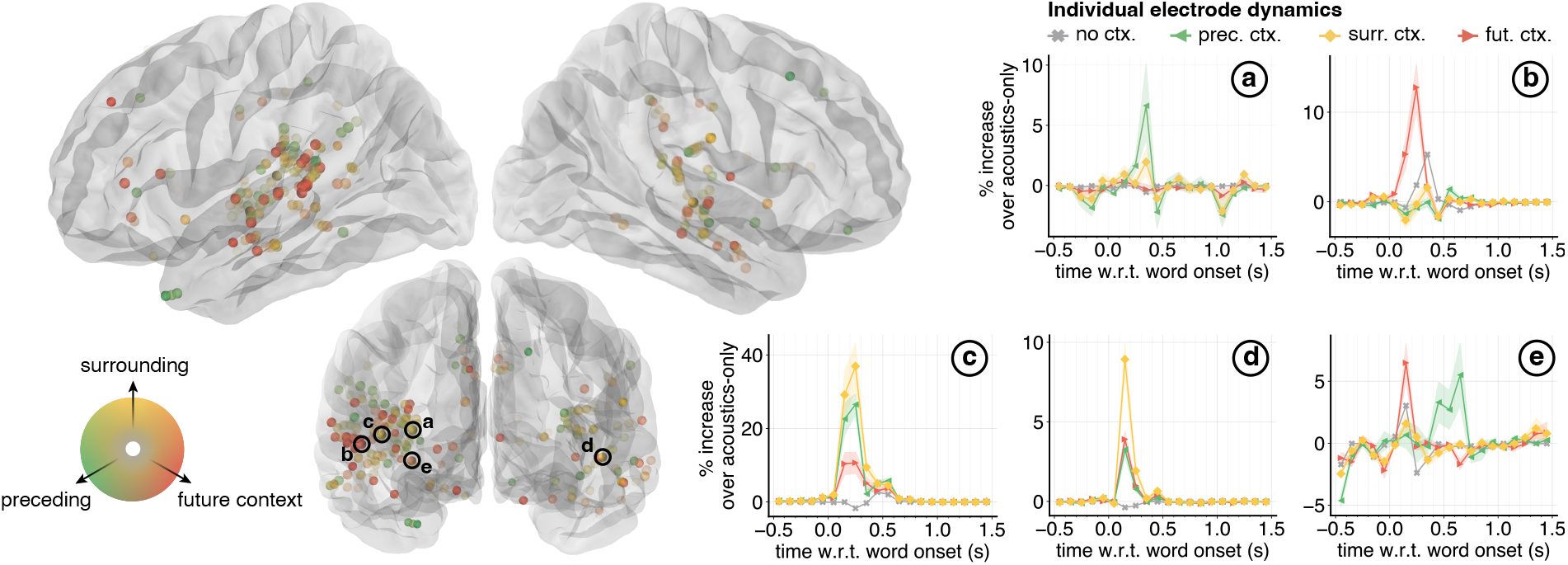
Electrodes vary in their Wav2Vec2-alignment to different context types. Electrodes are colored by triangulating the ratios of context-advantage scores for each context type, aggregating over scores between 0 and 300 ms after word onset. Shading in the epoch plots shows variation across context sizes within each type (std. err.).

Moreover, context preferences are spatially intermixed: electrode preferences do not cluster around particular anatomical subregions. Analyzing scores for a subset of 73 electrodes in auditory cortex (including only superiortemporal and transversetemporal electrodes) confirms that different subpopulations of electrodes are best modelled by different context types (Figure 6). Interestingly, surrounding context seems to best model a relatively smaller set of electrodes across the epoch, while it does show numerically strongest alignment with the neural activity that those electrodes record. This suggests that neural word representations supported by surrounding context are focal and strong, whereas future- and past-informed representations are more distributed and potentially more weakly encoded by individual populations.

**Figure 6:**
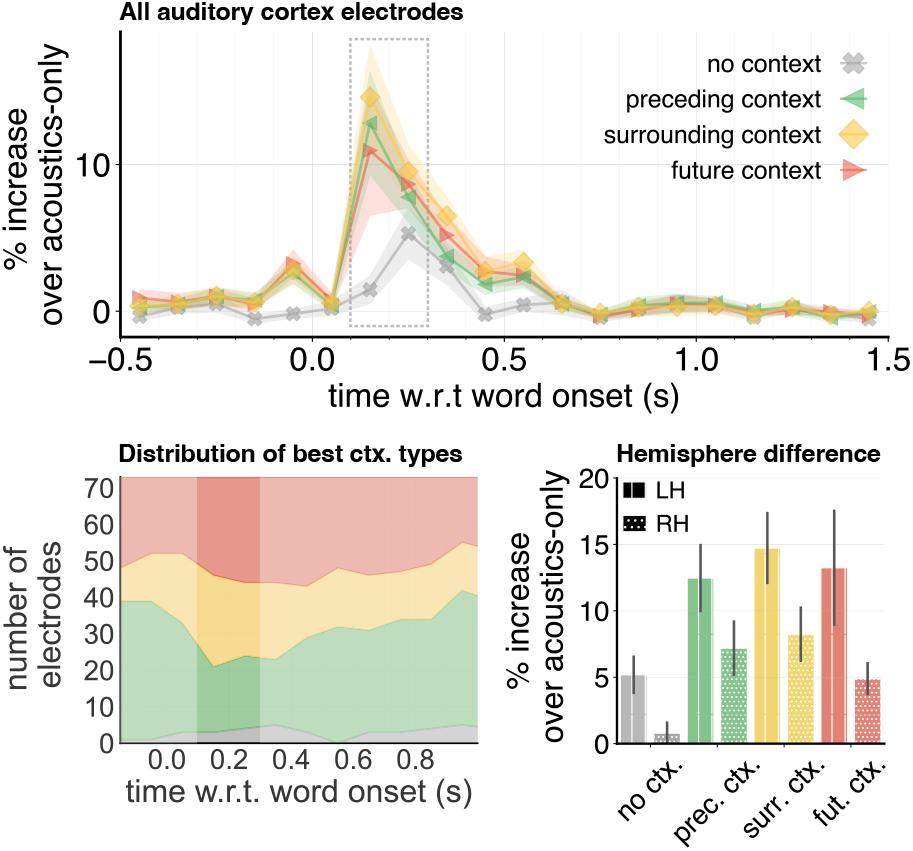
Among auditory cortex electrodes, there is no strong population-level preference for a specific context type, though context does improve neural alignment for almost all electrodes. In general, Wav2Vec2-alignment is higher in the left hemisphere than in the right.

Finally, the temporal resolution of iEEG data allows us to explore an open question that is unanswered by analyses on more temporally smeared signals such as fMRI: does context actually benefit neural alignment on the level of individual words, or does it facilitate longer time-scale representations that unfold across multiple words and phrases? The temporal dynamics of Wav2Vec2-alignment across the epoch suggest that contextual information boosts alignment during *current* word processing (i.e. within a relatively short time window after word onset); alignment therefore indeed seems to be driven by shared contextualized representations, rather than by neural responses to information preceding or following the target word.

## 6 Conclusions & Discussion

We have shown that including beyond-word context significantly affects the internal representational structuring of Wav2Vec2, affecting both its encoding of interpretable word-level features and its alignment to human intracranially-recorded neural activity in speech comprehension.

While it might be unsurprising that Wav2Vec2’s representational quality improves when processing inputs more similar to its training distribution, this insight is relevant for downstream applications of self-supervised speech representations where the encoding of linguistic information could be important. Previous studies interpreting self-supervised speech models (Choi et al., 2024) as well as their neural alignment (Oota et al., 2024) have noted the limited internal encoding of linguistic structure in S3Ms, but also mostly relied on alternative embedding extraction pipelines, e.g., using audio-sliced inputs with relatively little (or no) beyond-word context, or audio frame embeddings sampled without word alignment. Our results show that such embedding extraction decisions can significantly limit the internal representational capacity of S3Ms, potentially explaining part of the observed results.

To assess the neural alignment of context-informed Wav2Vec2 representations, we focused on Wav2Vec2’s unique contribution in modelling representation spaces in the human brain, over and above lower-level acoustic features. Earlier work predicting fMRI signals recorded during speech comprehension found earlier layers of Wav2Vec2 were best aligned to early speech processing regions (Millet et al., 2022), whereas we found highest model-to-brain alignment scores for Transformer layer 7 (Appendix B). We expect this to be a consequence of our strict control of low-level acoustics – earlier layers of Wav2Vec2 strongly encode acoustics (Figure 2), suggesting that it is important to control for acoustic correlates to enable interpretation of higher order model representations. When controlling for acoustics, we found that a substantial number of electrodes were better modelled when including context-informed Wav2Vec2 embeddings from layer 7. These representations align with dynamic neural activity in the temporal lobe, including the superior temporal gyrus and middle temporal gyrus, which are associated with phonemic, morphological, and lexical level processing (Hickok and Poeppel, 2007; Lu et al., 2026). Layer 7 is also where our representational probes for higher-order linguistic structures peak (Figure 2). We therefore hypothesize that relative beyond-acoustics alignment to electrode activity is *driven* by the encoding of higher-level linguistic structure in Wav2Vec2, though causal tests (e.g. using feature removal or disentanglement interventions on Wav2Vec2 embeddings, Krishnan et al., 2024; Mohebbi et al., 2024) would be crucial to properly test this hypothesis.

Comparing the relative neural alignment advantages between context types, we find variation across electrodes in the context type (preceding, surrounding, or future) that best modelled neural activity. The temporal dynamics of alignment to context-informed embeddings (Figures 5, 6) do not indicate that alignment to future or past context is driven by the presentation of future or past input; rather, different types of context support different brain-aligned representations of the target word. It is particularly intriguing that alignment with neural responses to the target word (within ≤ 300 ms after word onset) can be improved by providing (up to 20) *future context words* to the speech model. These effects cannot be explained by the brain’s processing of future words (as the neural response is measured before they occur); instead, they may reflect the prediction of upcoming speech content, or other representational links between the target word and its future context. Such mechanistic hypotheses should be investigated in future work.

The spatial intermixing of context type preferences echoes existing findings analyzing intracranial recordings in speech comprehension, showing that many different speech features are encoded in the temporal lobe (Yi et al., 2019). This suggests that contextual information is integrated within a recurrent circuit in the temporal lobe that leverages locally multiplexed representations of different context types (Gwilliams et al., 2025a). This account is inconsistent with models of human speech comprehension that posit discrete processing modules for different types of context, which would instead predict a primarily feedforward architecture.

Finally, we find higher unique Wav2Vec2-alignment in the left hemisphere as compared to the right, within auditory cortex (Figure 6). While lesion-symptom mapping clearly demonstrates left lateralized language function (Bates et al., 2003), intracranial studies often find symmetric encoding of speech and language features across hemispheres (Zhang et al., 2026; Bhaya-Grossman and Chang, 2022; Yi et al., 2019). The asymmetry we observe may again be due to controlling for acoustic variance when analyzing the neural alignment of S3M representations: it is plausible that the higher-level linguistic information encoded in Wav2Vec2 is what drives brain-to-model alignment when acoustic features are controlled for, and that this is prioritized by the left hemisphere.

To conclude, we demonstrated that the integration of beyond-word context is crucial both for Wav2Vec2’s word-level linguistic feature encoding, as well as its alignment to time-resolved human neural activity in speech comprehension. In the latter, we have shown promising benefits of controlling for low-level acoustics: we observe that neural activity in speech regions aligns with context-informed Wav2Vec2 representations from higher model layers, which also exhibit linguistic structuring. This emphasizes the potential of using self-supervised speech representations in modelling human spoken language processing, not only for replicating known variation along the cortical hierarchy, but also for generating hypotheses on the kinds of neural representations involved in the linguistic interpretation of speech signals. In particular, our finding that spatially neighboring yet distinct neural populations in the temporal lobe encode representations shaped by different contextual sources (past, future, and surrounding input) provides key insight into the neural circuitry that integrates multiple forms of contextual information, to ultimately derive speech comprehension.

## 7 Limitations

We have reported findings based on one 15 minute single-speaker recording; all our analyses therefore capture within-speaker variation only. Speaker effects on Wav2Vec2’s encoding of linguistic information, as well as the generalizability of such information across speakers, remain to be explored. This also holds for potential effects of speaker information on Wav2Vec2’s alignment to neural activity.

Using RSA to measure the alignment of Wav2Vec2 representations to brain activity diverges from existing work testing how well fMRI signals can be predicted from Transformer hidden states using ridge regression (e.g. Millet et al., 2022; Vaidya et al., 2022). By fitting only one weight per included feature space instead of separate weights for each feature dimension, our weighted representational model includes fewer degrees of freedom in the mapping from feature to neural representation space. Although it is exciting that we still observe model-brain alignment using such a restrictive mapping approach, it is unclear from our main analyses whether our observed context effects would still hold when more complex mappings could be learned (e.g. by optimizing weights for individual feature dimensions, as in earlier work). To verify the relevance of our findings to more commonly used approaches for measuring the neural predictivity of model representations, we include results using ridge regression instead of RSA in Appendix C. These results replicate the general advantage of context-informed over context-uninformed embeddings, as well as observed differences between context types.

In manipulating the amount of context available to Wav2Vec2, we silence-padded all audio segments to be the same duration, to avoid confounding effects of added context with effects of increased input duration (subsection 3.2). Audio segments with relatively little speech content and long silences are potentially out of distribution for Wav2Vec2 compared to what it has been trained on, which could be an additional reason for the decreased representational quality of context-uninformed embeddings. To inspect effects of silence padding on our results, we repeated our main analyses using embeddings extracted without silence padding (i.e. audio-slicing inputs around the included context windows without adding silence); results are included in Appendix D and generally look very similar to our main results with silence-padded embeddings.

For our neural alignment analyses, we report the electrode-wise changes in RSA score when including Wav2Vec2 features in our representational model, quantified as the percentage increase compared to scores obtained with only low-level acoustic features (envelope+MFCC). While this gives a good view of the relative benefit of Wav2Vec2 features for each electrode, percentage rescaling also somewhat obscures variance in absolute RSA scores across electrodes. Variance in absolute RSA scores can arise from variation between participants or anatomical regions, as well as electrode-wise variation in signal quality; we have not further investigated such variation here. We refer to supplementary figures S10, S8 and S7 for an idea of the absolute RSA score vs. percentage increase distributions. We generally observe the same context-advantage patterns when analyzing differences in terms of absolute RSA scores instead of percentages (Figure S9), as well as differences in absolute ridge regression scores (Figure S11).

The participants in our study were patients undergoing treatment for drug resistant epilepsy; thus, all participants have a clinically diagnosed neurological disorder. We mitigate concerns this may have regarding the generalisation of our results by only including data from patients who had no clinical history of speech or language processing deficits, and by only including electrode contacts in our analyses that were outside of the clinically determined pathological tissue. Future work should aim to replicate these findings in neurologically healthy individuals using time-resolved non-invasive neural recording techniques.

Finally, our analyses focused on a subset of electrodes mostly concentrated in early auditory and speech processing regions (Figure 4); our coverage of anatomical regions is limited by the sparse localisation of sEEG electrodes across the brain (Figure 1). It is likely that context also affects Wav2Vec2-alignment to other brain regions that we were unable to sample in this study. The broader anatomical distribution of such effects remains to be explored.

## 8 Ethical considerations

All stereoelectroencephalography (sEEG) data were acquired from patients undergoing clinically indicated monitoring for drug-resistant epilepsy; no electrodes were implanted for research purposes. The study was approved by the local institutional review board, and participants provided informed consent to take part in this research study. Consent emphasized voluntariness, the ability to withdraw at any time, lack of direct clinical benefit, and scheduling that does not interfere with standard care (Chiong et al., 2018).

## 9 Acknowledgements

MdHK is funded by the Netherlands Organization for Scientific Research (NWO), through Gravitation Grant 024.001.006 to the Language in Interaction Consortium. LG is funded by the Esther A. & Joseph Klingenstein Fund, Whitehall Foundation 2024-08-043, and BRAIN Foundation A-0741551370.

## A Representational probing details

### A.1 Acoustic alignment

To quantify the layer-wise acoustic structuring of Wav2Vec2 word embeddings, we compute alignment to 13-dimensional MFCC features extracted for all 2001 words in the narrative, mean pooled across each word duration. The acoustic *MFCC-CCA score* is the mean Pearson’s correlation between the 12 components projected from Wav2Vec2 and MFCC space for each word in the test split, after fitting the CCA model on the train split.

### A.2 Syllable type clustering

To get a layer-wise estimate of word form structuring in Wav2Vec2 embeddings, we probe a subset of 108 mono- and disyllabic words for their consonant-vowel (syllable type) structure. The subset spans 6 syllable types (CVC, CV, CVCC, VC, CVCV, CVCVC) for which enough unique word occurrences could be sampled from the narrative data; we ensured that multiple unique words were included for each syllable type and that words did not overlap between train and test splits, to avoid confounding of word identity. The *SylType-LDA* score is the silhouette score (Rousseeuw, 1987) quantifying clustering by syllable type in the 5-dimensional LDA projections of test words, after fitting the LDA model on words in the train split. Silhouette scores range between −1 and 1, where 0 corresponds to random structure and positive values indicate more well-separated clusters.

#### A.3 Semantic alignment

To quantify alignment with semantic structure in Wav2Vec2 embeddings, we probe a subset of 463 unique words from 3 part-of-speech categories (212 nouns, 162 verbs, 89 adjectives) and use CCA to measure the alignment of their Wav2Vec2 embeddings to static text-based word embedding vectors designed to capture lexical semantic structure (GloVe; Pennington et al., 2014). We evaluate semantic alignment separately within each POS category and ensure that test samples consist exclusively of unique word forms, to avoid confounding of word identity and part-of-speech information. The semantic *GloVe-CCA score* is the mean Pearson’s correlation between the 12 components projected from Wav2Vec2 and GloVe space for each word in the test split, after fitting the CCA model on the train split.

### A.4 Part-of-speech clustering

To measure the degree of structuring by syntactic word category in Wav2Vec2 embeddings, we sample equal-sized sets of word *token* occurrences from a total of 205 unique word *types*, spanning 6 part-of-speech categories (55 nouns, 52 verbs, 31 adverbs, 21 adjectives, 18 adpositions, 28 pronouns), using 100 token occurrences of each POS category for training and 20 token occurrences for testing. We ensure that each test set contains exactly 2 token occurrences of 10 word types which are not used in training, to avoid effects of word identity. The *POS-LDA* score is the silhouette score (Rousseeuw, 1987) quantifying clustering by part-of-speech in the 5-dimensional LDA projections of test words, after fitting the LDA model on words in the train split. Silhouette scores range between −1 and 1, where 0 corresponds to random structure and positive values indicate more well-separated clusters.

### A.5 Supplementary results

In Figure 2 we visualized probing results for embeddings extracted from a Wav2Vec2 model pretrained on 960 hours of speech recordings (Baevski et al., 2020). Here we verify that the information we probed for can be extracted from model embeddings as a result of model training, rather than resulting from a propagation of input acoustics throughout model layers that could already be present in an untrained model (Chrupała et al., 2020). Figures S1, S2, S3 and S4 visualize results for each probe task and context condition individually, including scores for a randomly initialized Wav2Vec2 architecture that has not been trained. These results confirm that the observed layerwise probing results are a consequence of model training: results on untrained model embeddings generally show no layerwise developments, and are always lower than scores obtained for trained model embeddings at their layerwise peaks. Moreover, for any task probing linguistic structure beyond acoustics, only some layers from the trained Wav2Vec2 model show consistent above-random performance.

In Figure S5, we additionally include a visual demonstration of the observed layerwise differences in our clustering probes, by plotting test item projections of our POS-LDA probe along the first three (out of five) extracted LDA directions.

**Figure S1:**
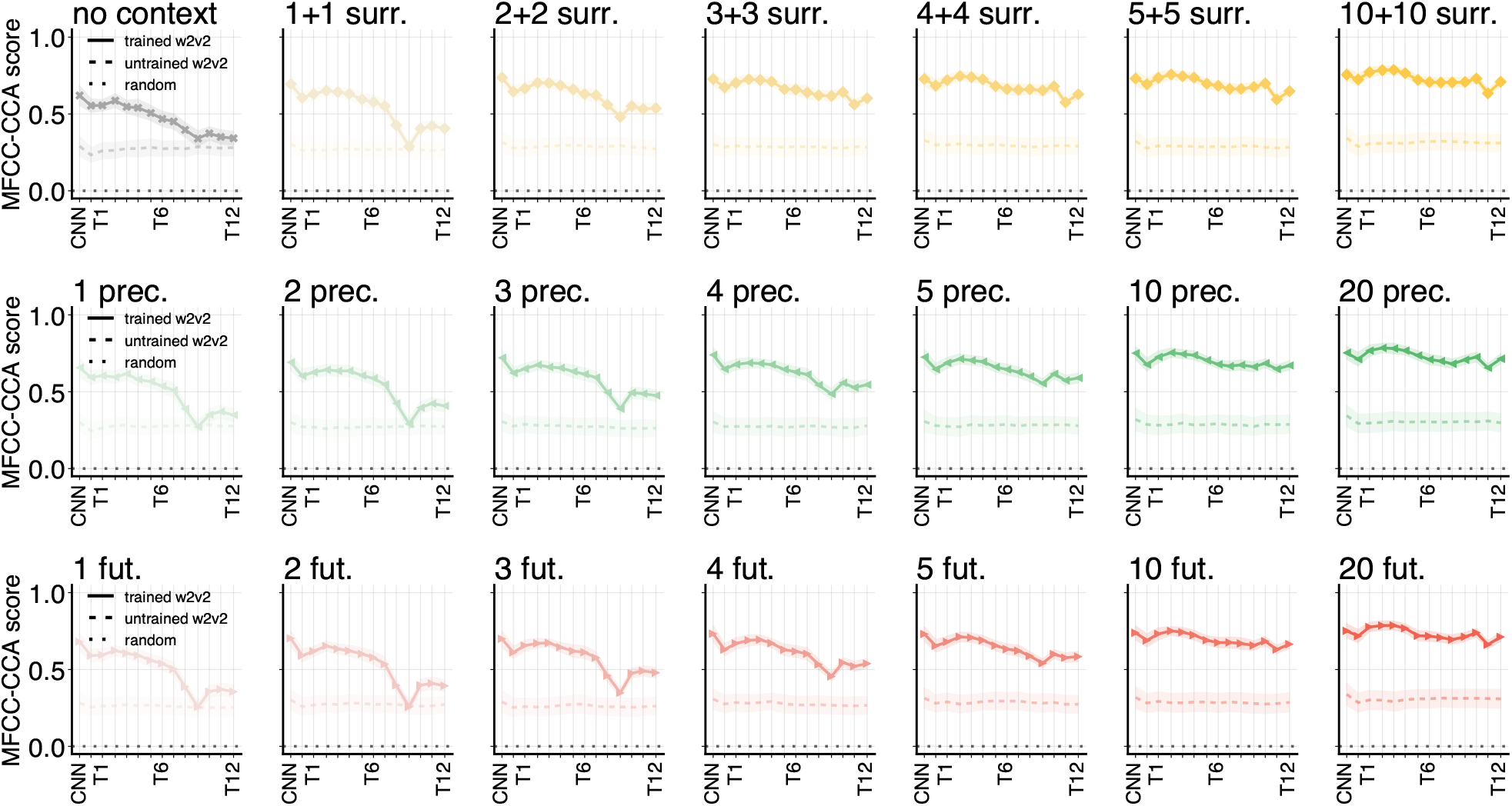
Layerwise **acoustic CCA** results for all context conditions and model layers, including scores for trained (solid line) and untrained (dashed line) Wav2Vec2 models. Black dotted lines mark scores indicating random structure; shading visualizes variance between different test splits (standard error).

**Figure S2:**
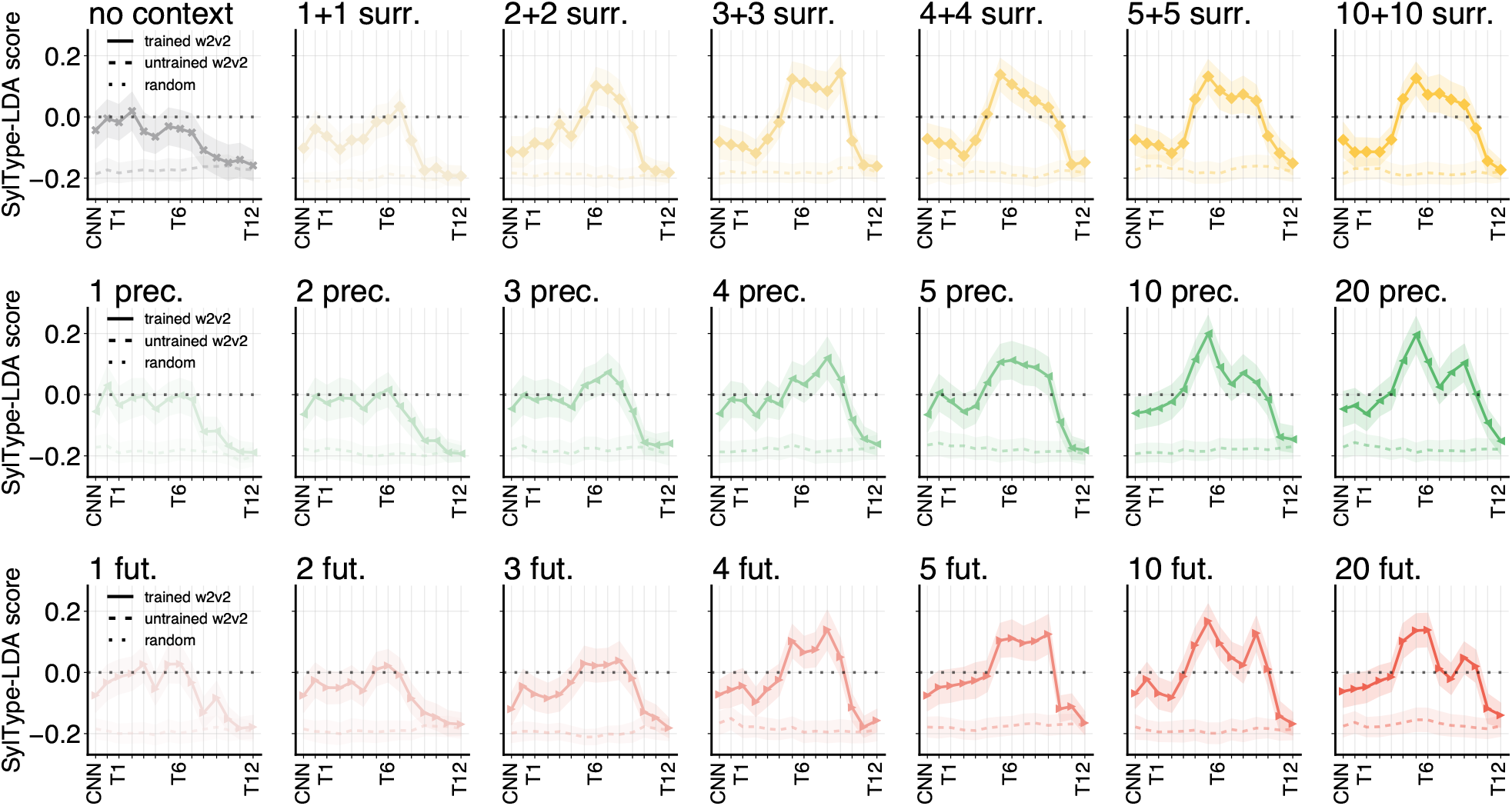
Layerwise **syllable type LDA** results for all context conditions and model layers, including scores for trained (solid line) and untrained (dashed line) Wav2Vec2 models. Black dotted lines mark scores indicating random structure; shading visualizes variance between different test splits (standard error).

**Figure S3:**
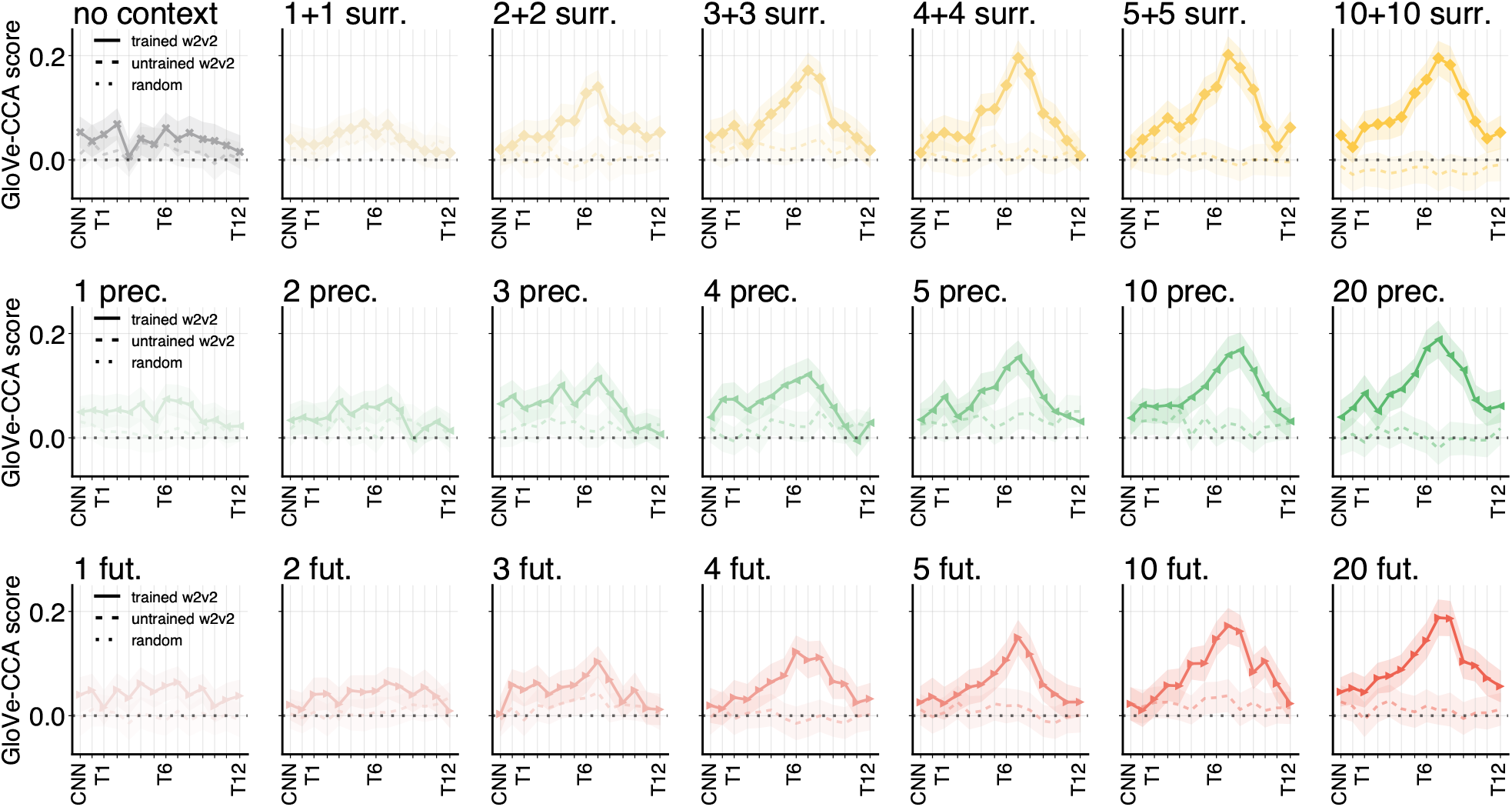
Layerwise **semantic CCA** results for all context conditions and model layers, including scores for trained (solid line) and untrained (dashed line) Wav2Vec2 models. Black dotted lines mark scores indicating random structure; shading visualizes variance between different test splits (standard error).

**Figure S4:**
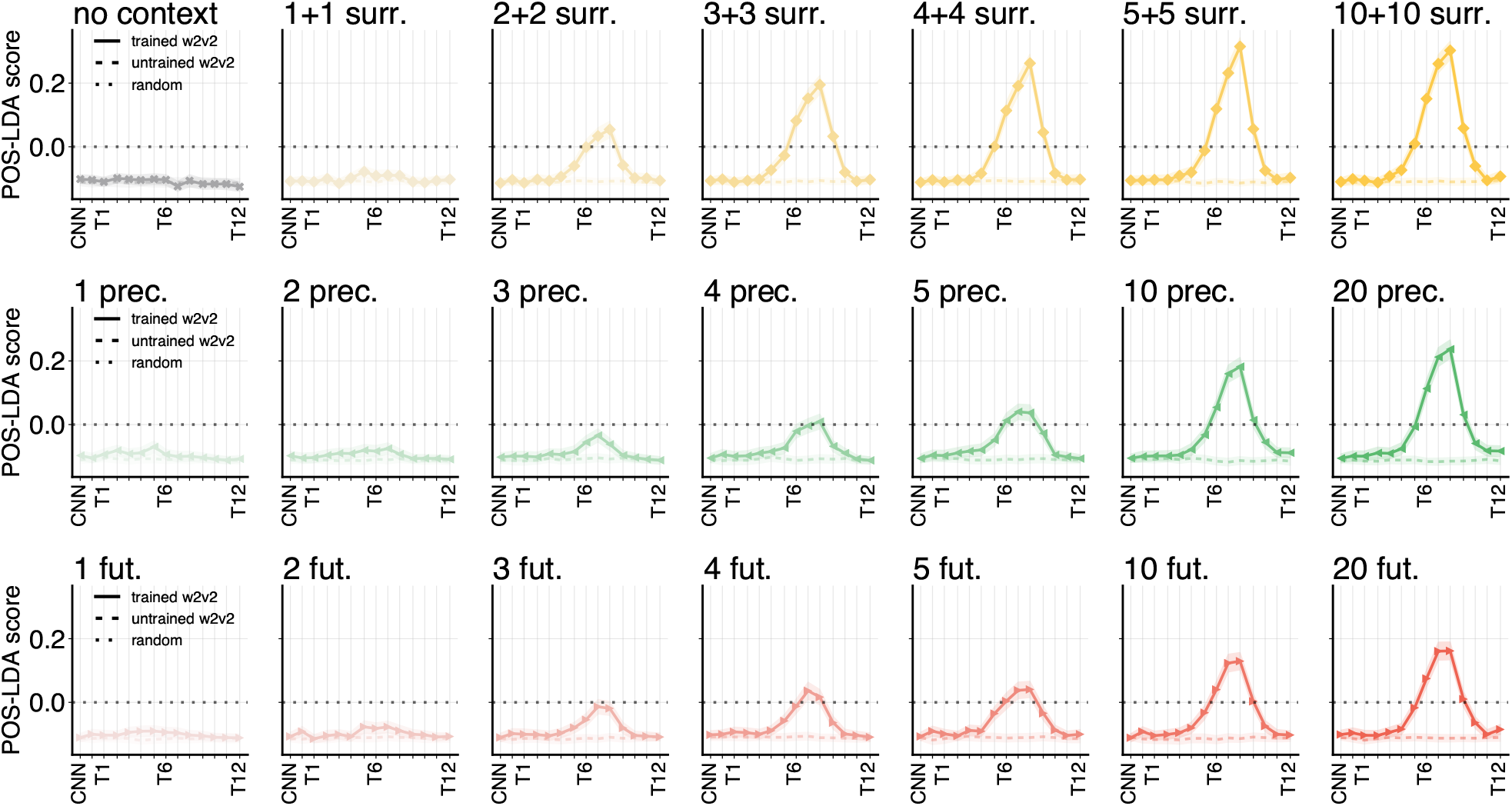
Layerwise **part-of-speech LDA** results for all context conditions and model layers, including scores for trained (solid line) and untrained (dashed line) Wav2Vec2 models. Black dotted lines mark scores indicating random structure; shading visualizes variance between different test splits (standard error).

**Figure S5:**
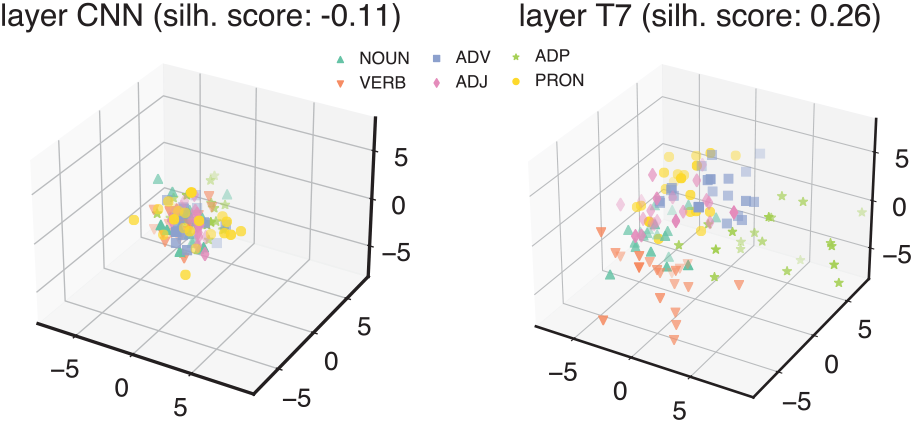
Comparing POS projections extracted from two layers of the trained Wav2Vec2 model in the same context condition (including 10+10 surrounding words), we verify that the measured layerwise differences in silhouette score indeed capture observable differences in clustering by POS category across model layers. Data points are projections for individual words in the POS probe test set, colored by their POS category. We note that silhouette scores are computed across all 5 discriminative directions projected by the LDA probe, and we here only visualize 3.

## B Electrode selection and neural alignment details

### B.1 Model structure

For each context-informed embedding condition *c* and each epoch time window *t*, our NNLS model is structured as follows:

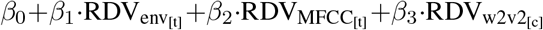

i.e., as a linearly weighted sum over representational dissimilarity vectors (RDVs) from three feature spaces: two defined by low-level acoustic features local to time window *t*, and one defined by the word-level Wav2Vec2 representations extracted in context condition *c*. The RDVs contain the pairwise cosine distances within each normalized feature space, between all 2001 words in the spoken narrative (for the acoustic features: between the local acoustics of each time window *t* relative to word onset). As local acoustic features, we intend to capture both temporal and spectral variation in the speech signal at each time window, by including RDVs constructed based on the *speech envelope* (containing distances between envelope vectors of 10 concatenated values sampled across each time window *t*), and based on *MFCC* features (containing distances between 13-dimensional vectors mean-pooled across each time window *t*). For Wav2Vec2, we use the same word-level RDV across all time windows in the epoch, expecting to measure contributions of Wav2Vec2 to occur only at time windows where shared beyond-acoustic information related to target words is encoded.

### B.2 Model evaluation

We fit RDV models to optimize predictions of the neural RDV corresponding to the word-locked iEEG signal at time window *t* (i.e.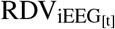). We evaluate our RDV models by 3-fold cross-validation, training each NNLS model on two blocks of the narrative and testing on the remaining third. As our *RSA score* measure for evaluating neural representational similarity, we compute the Pearson’s correlation between the combined feature RDV and the neural RDV for each fold. To obtain the *unique contribution* of Wav2Vec2 to neural representational similarity, we apply variance partitioning (Lescroart et al., 2017; de Heer et al., 2017), comparing scores obtained with our full RDV model (including 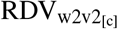) with an alternative RDV model including only low-level acoustics (excluding 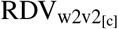). For all our main analyses, we thus report unique *Wav2Vec2-alignment* scores, defined as the *relative change* in neural similarity when including Wav2Vec2, computed as a percentage of the score obtained with the acoustics-only model.

### B.3 Quantifying context-preference

We aim to explore to what extent electrode activity is modelled better with context-informed word embeddings vs. non-contextual embeddings, and also to what extent different electrodes or time windows benefit from particular context *types* (past, surrounding, or future).

With this purpose in mind, we quantify the unique benefit of each context condition to neural representational similarity, the *context-advantage score*, as the difference in Wav2Vec2-alignment between each context-including condition and the no-context condition. We then define a *context-preference* score for each context type, by averaging context-advantage scores over all contexts within each type, and computing preference as the fraction of each type’s mean context-advantage in the sum of all positive mean context-advantages across the three types. In other words, electrode-level preference scores are defined for each context type, ranging from 0.33 (if context-advantage scores are equal for all context types) to 1 (if only one context type shows an advantage over the no-context condition), and only for electrode time windows where at least one context-informed condition proves beneficial over the no-context condition.

### B.4 Supplementary results

In Figures S6, S8, S7, S9, and S10, we visualize results that complement the findings we report in subsection 5.3. Relative Wav2Vec2-alignment peaks in layer 7 of the trained Wav2Vec2 model when controlling for low-level acoustics, and is substantially higher for any context-informed embedding than for isolated word embeddings (Figure S6). Wav2Vec2 features consistently improve alignment of our representational model to intracranially-recorded neural activity, as demonstrated by the distribution of relative % improvement scores across electrodes (Figure S7), as well as the positive shift in electrode-level RSA scores comparing the distributions of absolute acoustics-only vs. full model scores (Figure S8).

Context effects on neural alignment are similar when analyzing absolute differences in RSA scores without percentage rescaling (Figure S9). Mean absolute contributions of untrained Wav2Vec2 embeddings are negative, whereas their mean percentage improvement scores are positive (Figure 4) – this is an effect of percentage rescaling: when the contribution of untrained Wav2Vec2 embeddings is negative for electrodes with relatively high acoustics-only RSA scores, the *percentage decrease* driven by Wav2Vec2 features in these electrodes may be relatively small, while the *decrease in absolute score* is relatively high, compared to other electrodes.

**Figure S6:**
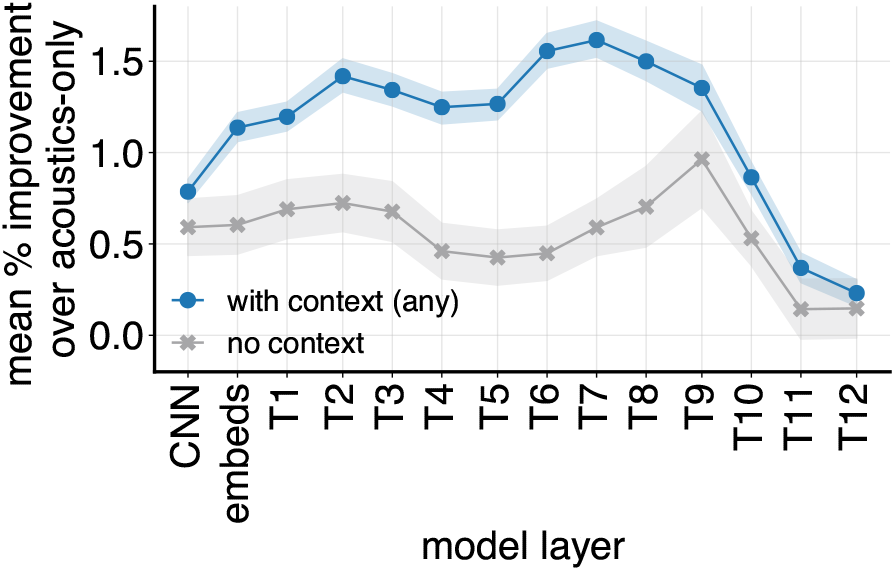
Unique Wav2Vec2-alignment scores across model layers, defined as the relative % increase over the acoustics-only model. We visualize mean scores across electrodes and epoch time windows.

**Figure S7:**
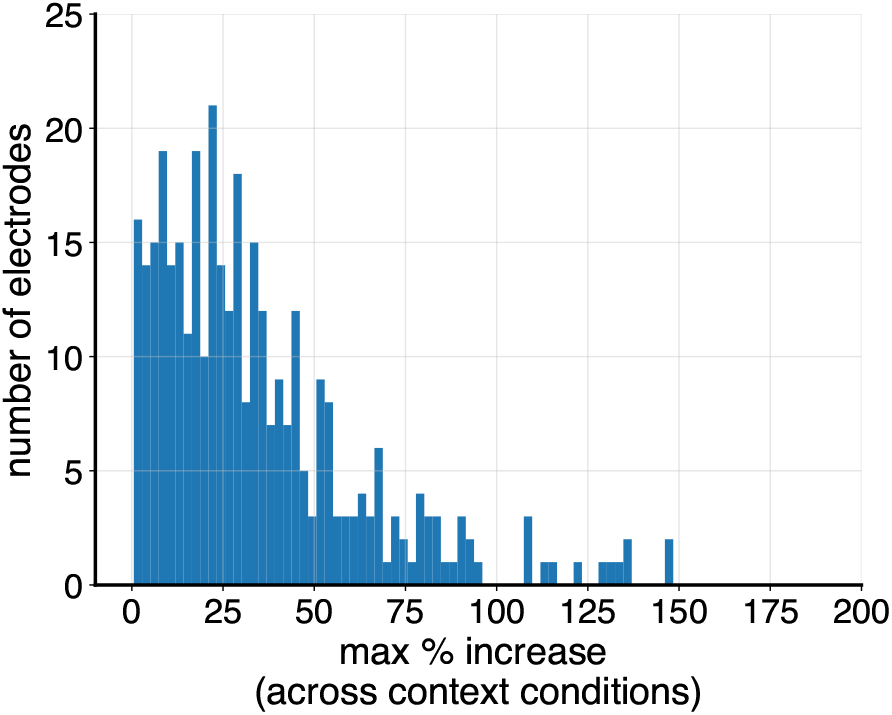
Distribution of unique Wav2Vec2-alignment scores quantified as the relative % increase over the acoustics-only model. We here include the maximum score across the epoch for each electrode.

**Figure S8:**
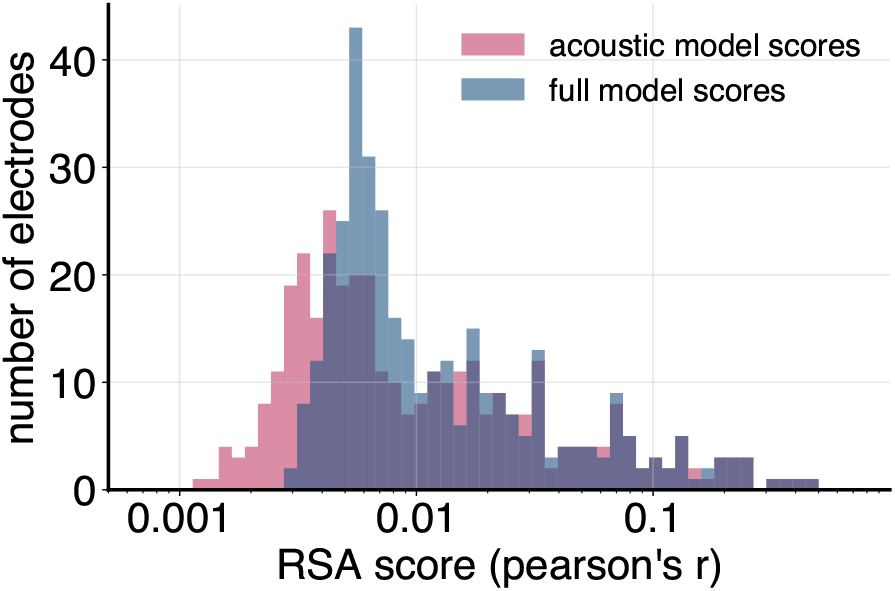
Distribution of acoustics-only vs. full model scores (pearson’s r correlation between the feature and neural RDVs). We include the maximum score across the epoch for each electrode. The x-axis scale is logarithmic. Full model scores (which include Wav2Vec2 features) are consistently higher than acoustics-only scores.

**Figure S9:**
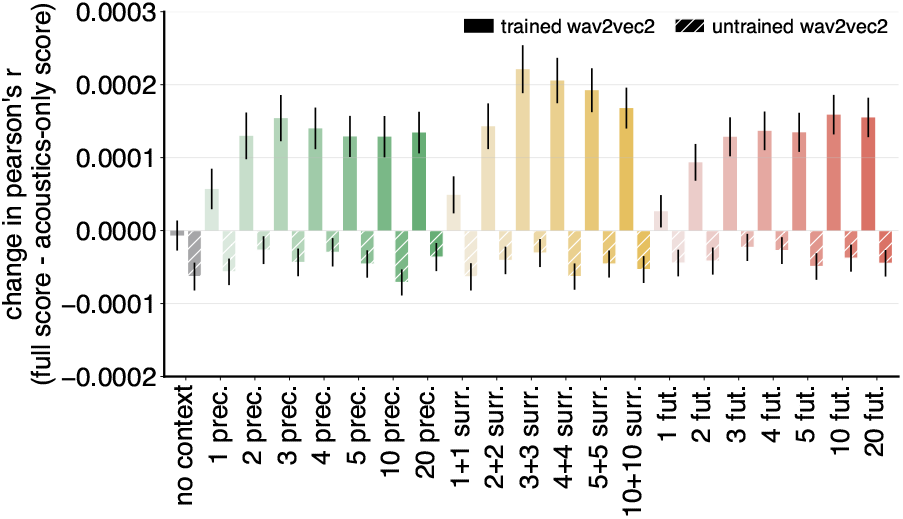
The pattern of context effects on neural alignment is similar when analyzing absolute changes to RSA scores instead of % improvement scores (compare Figure 4).

**Figure S10:**
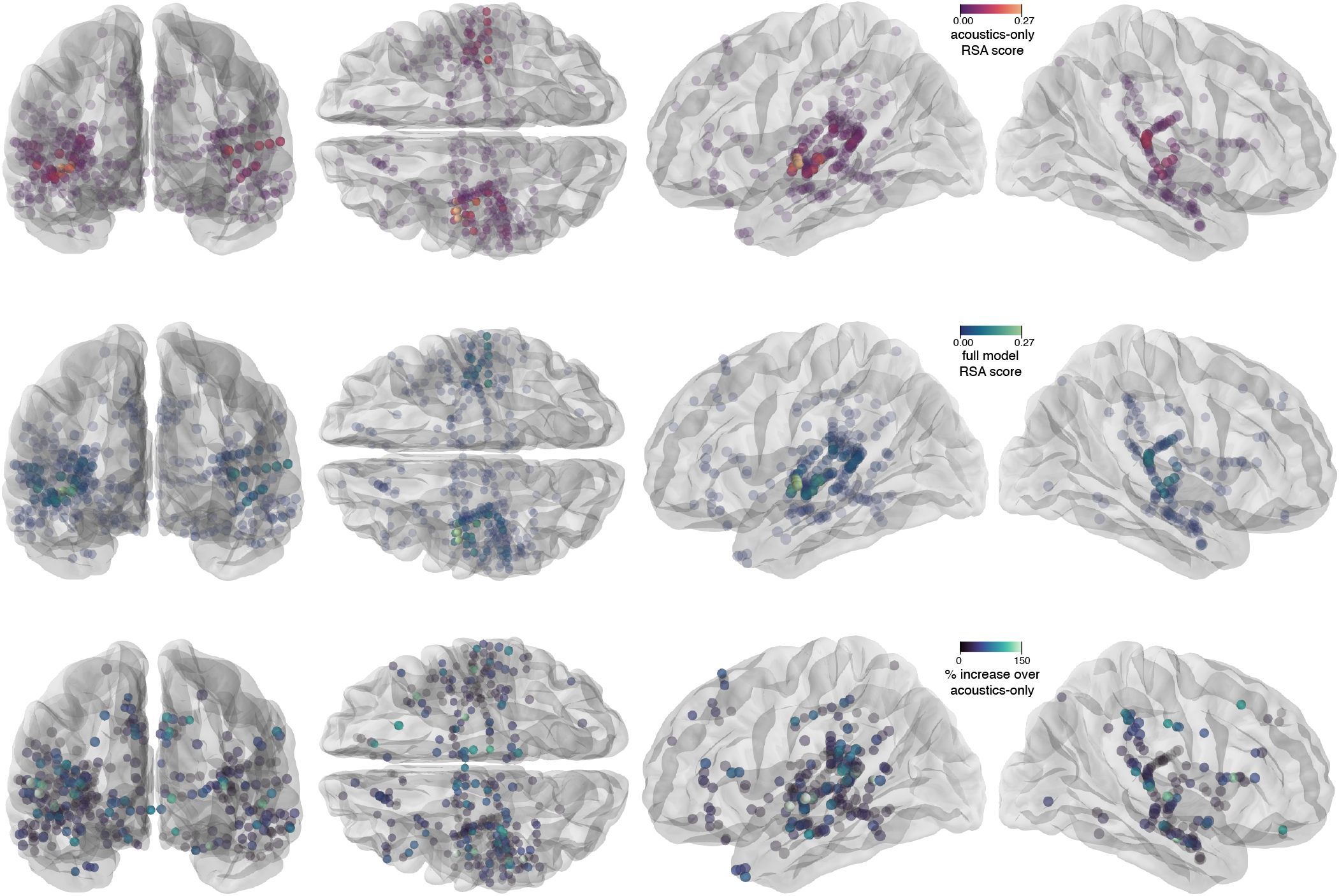
We visualize the spatial distribution of model scores across electrodes, from 4 different views of the brain (caudal, dorsal, left- and right-lateral). The visualized scores are the maximum scores between 0 and 300 ms after word onset. Top row: RSA scores obtained with models including only acoustic features (envelope+MFCC). Middle row: RSA scores obtained with full models (envelope+MFCC+Wav2Vec2). Bottom row: relative electrode-wise % increase of the full model over the acoustics-only scores. (a) Distribution of acoustics-only vs. full model scores obtained with ridge regression (pearson’s r correlation between the predicted and observed neural signal). We include the maximum score across the epoch for each electrode. The x-axis scale is logarithmic. Correlations between ridge-predicted and observed neural activity are generally higher than those between representation space (dis)similarities (compare Figure S8), but we observe a similar benefit of including Wav2Vec2 on top of acoustics-only features. (b) For scores obtained with ridge regression, we observe that the contribution of Wav2Vec2 is negative across most of the epoch, but positive within a short time window after word onset (*±*0 − 300 ms). (c) For ridge scores within 0 − 300 ms after word onset, we see a clear advantage of context-informed over context-uninformed embeddings, as well as a benefit for the surrounding context type (similar to scores obtained with RSA; see Figures 4, S9). As for context size, more context seems to generally improve scores obtained with ridge regression, somewhat diverging from patterns observed with RSA.

## C Ridge regression vs. RSA

Our approach to neural-representational modelling (Appendix B) predicted distances in neural representation space as a weighted sum of distances in several feature representation spaces. While this variant of RSA has been explored for measuring neural alignment in the domain of vision (Khaligh-Razavi and Kriegeskorte, 2014; Jozwik et al., 2016), we are not aware of studies in the speech domain using a similar approach.

In this section we explore whether our main results replicate when directly predicting recorded neural activity itself, more in line with earlier work predicting fMRI signals using self-supervised speech model representations (Millet et al., 2022; Vaidya et al., 2022). Following these earlier studies, we fit an 𝓁2-penalised linear regression (ridge regression) model to predict neural activity *Y*_*train*_ from feature set *X*_*train*_, and then evaluate predictions of unseen neural activity against true measurements by computing the Pearson’s correlation between predicted *Ŷ* _*test*_ and true *Y*_*test*_. We otherwise replicate our RSA set-up as closely as possible; cross-validating scores across three folds of the data, including both acoustic features and Wav2Vec2 representations in *X*, and analyzing our effects of interest on the unique contribution of Wav2Vec2. We fit and evaluate individual ridge regression models for each of the 21 context conditions and 100 timepoints across the epoch.

Effects of context on neural predictivity as evaluated using ridge regression generally follow those based on RSA reported in our main text (Figure S11). Correlations of ridge-predicted and observed neural activity are higher than the correlations observed in our RSA analyses, but do show similar benefits for context-informed Wav2Vec2 embeddings. One difference between the ridge- and RSA-based results is that ridge-based measurements indicate that improvements to neural alignment do not plateau or decrease at larger context sizes (compare Figure S11c and Figure S9). It is possible that more distant but neurally relevant contextual information is stored in a relatively small subspace of model embeddings, which can be extracted when fitting weights for individual embedding dimensions with ridge, but which is not well-represented in the cosine-based (dis)similarity spaces constructed for RSA. We leave further explorations of differences between neural alignment methodologies to future work.

**Figure S11:**
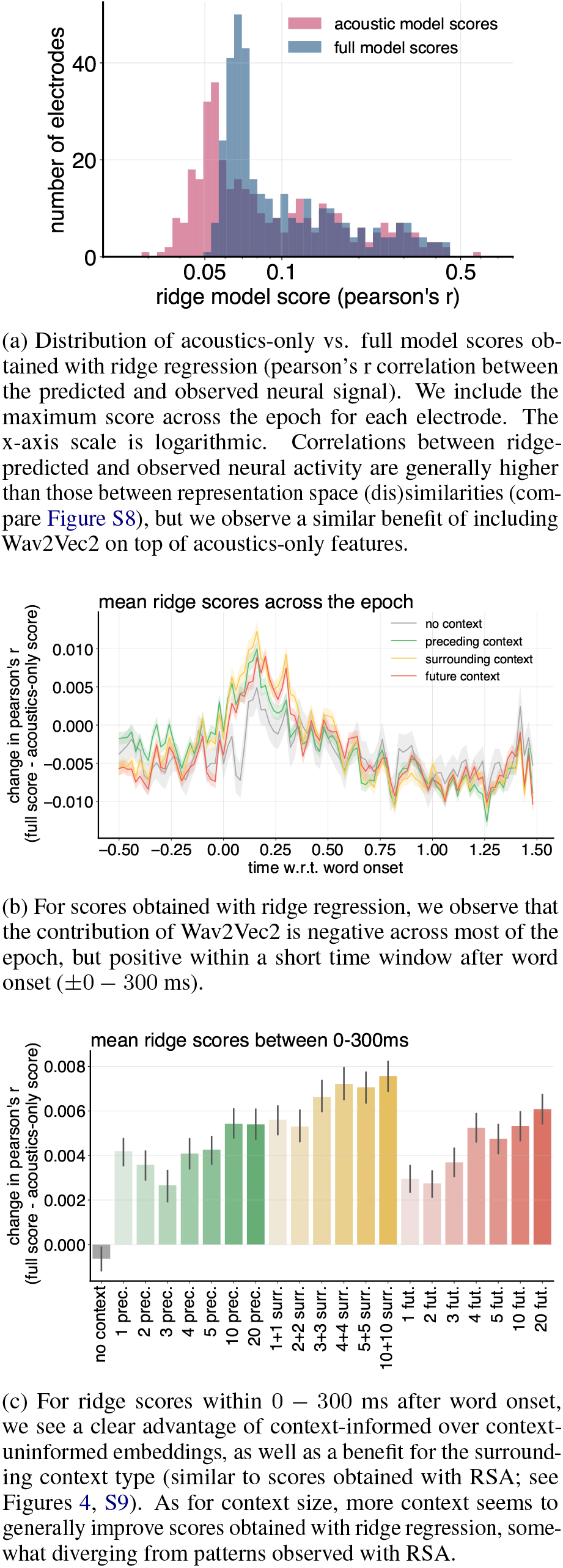
Our main results replicate when predicting neural activity using ridge regression instead of modeling (dis)similarities in neural representation spaces using NNLS.

## D Results without silence-padding

To confirm that our observed effects are not driven by the silence-padding of audio inputs, we repeated our main probing and neural alignment analyses using embeddings extracted without silence-padding. Results are visualized in Figure S12 and Figure S13 respectively and are generally very similar to those observed without silence-padding (compare Figures 2 and S9). We note two minor differences:

1. context effects on neural alignment are generally slightly smaller without silence-padding, with more negative scores for isolated word embeddings;
2. the peaks for our syllable type clustering probe are more concentrated in a single, higher, model layer. We leave further interpretation of these differences to future work.

**Figure S12:**
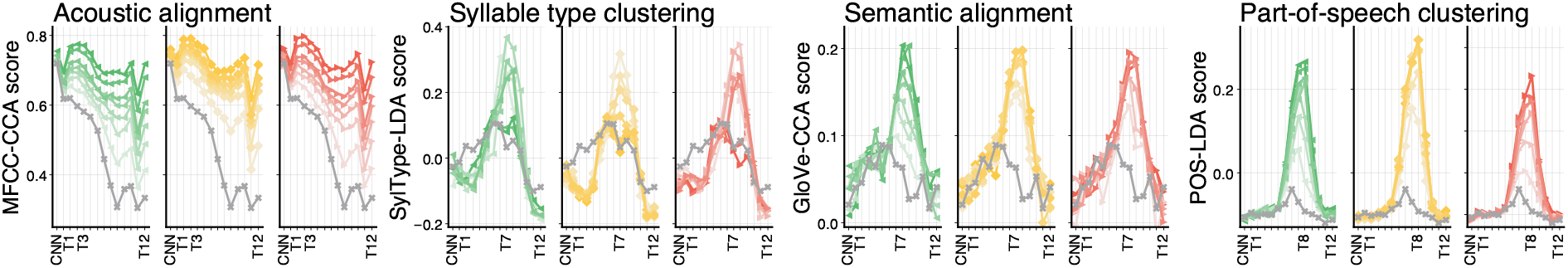
Probing scores for embeddings extracted without silence-padding show generally similar layerwise patterns to those observed for embeddings extracted with silence-padding (compare Figure 2).

**Figure S13:**
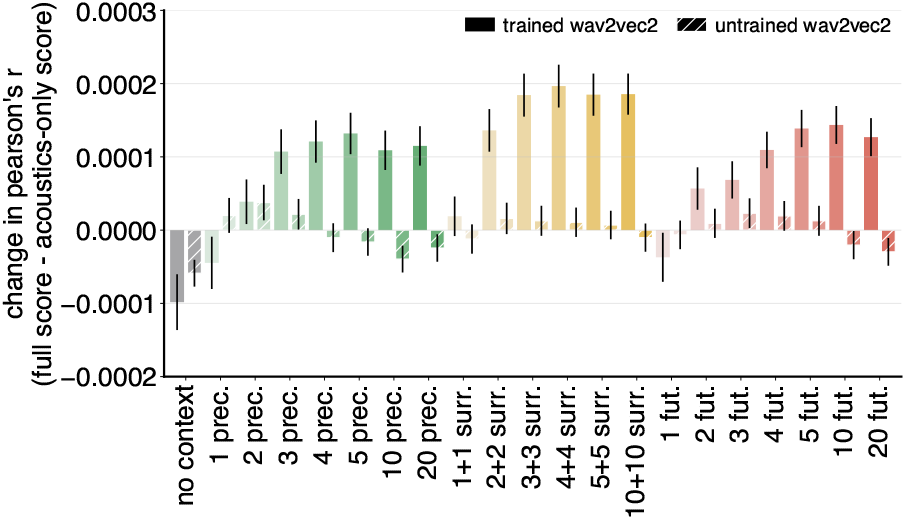
Context effects on neural alignment show generally similar patterns for embeddings extracted without silence-padding compared to those observed for embeddings extracted with silence-padding (compare Figure S9).

## Notes

### Competing Interest Statement

The authors have declared no competing interest.

